# Merqury: reference-free quality, completeness, and phasing assessment for genome assemblies

**DOI:** 10.1101/2020.03.15.992941

**Authors:** Arang Rhie, Brian P. Walenz, Sergey Koren, Adam M. Phillippy

**Affiliations:** Genome Informatics Section, Computational and Statistical Genomics Branch, National Human Genome Research Institute, National Institutes of Health, Bethesda, MD USA

**Keywords:** Genome assembly, assembly validation, benchmarking, k-mers, haplotype phasing, trio binning

## Abstract

Recent long-read assemblies often exceed the quality and completeness of available reference genomes, making validation challenging. Here we present Merqury, a novel tool for reference-free assembly evaluation based on efficient k-mer set operations. By comparing k-mers in a *de novo* assembly to those found in unassembled high-accuracy reads, Merqury estimates base-level accuracy and completeness. For trios, Merqury can also evaluate haplotype-specific accuracy, completeness, phase block continuity, and switch errors. Multiple visualizations, such as k-mer spectrum plots, can be generated for evaluation. We demonstrate on both human and plant genomes that Merqury is a fast and robust method for assembly validation.

**Availability of data and material:** Project name: Merqury

Project home page: https://github.com/marbl/merqury, https://github.com/marbl/meryl

Archived version: https://github.com/marbl/merqury/releases/tag/v1.0

Operating system(s): Platform independent

Programming language: C++, Java, Perl

Other requirements: gcc 4.8 or higher, java 1.6 or higher

License: Public domain (see https://github.com/marbl/merqury/blob/master/README.license) Any restrictions to use by non-academics: No restrictions applied

## Background

With recent advances in long-read^1–3^ and long-range sequencing technologies^4–6^, new assembly pipelines are generating more continuous, complete, and accurate diploid genome assemblies than ever before^4,7–14^.

However, *de novo* assembled genomes are difficult to validate due to the lack of a known truth. Existing methods use Illumina reads to infer base-level accuracy by aligning the reads to the assembly for evaluation^15^. Base errors in the consensus are detected as variants (SNPs or small indels) when aligning the short reads. However, this method is heavily reliant on the short-read mapping, which could be biased in repetitive regions, under-collapsed regions, or regions of low consensus accuracy. For measuring completeness and false duplications, near-universal single-copy orthologs (BUSCOs)^16^ have been widely used to evaluate the gene content of assemblies. BUSCO is robust for species that have been widely studied, such as human and mouse. However, this analysis can be inaccurate when the newly assembled genome contains true copy number or sequence variants that were not considered when building the initial BUSCO gene set. In addition, BUSCO only examines conserved single-copy genes and fails to evaluate the most difficult-to-assemble regions of the genome.

In contrast, k-mers (genomic substrings of length *k*) can be used in a reference-free manner for assessing genome assembly quality metrics. Genome assembly validation via k-mer copy number analysis was introduced by Mapleson *et al*. in their KAT tool^17^, which enables visual inspection of k-mer spectra plots to identify artificial duplications and missing sequences. Merqury takes much of its inspiration from the ideas introduced by KAT. QUAST-LG^18^ is another assembly validation tool that provides both BUSCO and KAT measurements, as well as alignment against a closely related reference genome. However, many of QUAST’s metrics are based on the reference alignment, which incorrectly reports true variants in the assembled genome as potential mis-assemblies.

Assessing haplotype phasing without a truth set is also challenging. Diploid genome assemblers generate both primary and alternate assemblies representing the two haplotypes. The primary assembly is typically a pseudo-haplotype that captures both the homozygous regions along with a single copy of the heterozygous alleles. Such a pseudo-haplotype does not guarantee long-range phasing, so to estimate phase block statistics, the alternate alleles must be mapped back to the primary assembly to determine regions corresponding to the primary-alternate haplotype phase blocks^9^. However, this can be challenging when the alternate alleles do not map well to the primary sequence due to high sequence divergence or mis-assemblies. Moreover, long-read assemblies often collapse regions of low heterozygosity, which are excluded when calculating phase block statistics, thus over-representing the correctness.

Alternative methods report phasing statistics from small variants (mostly SNPs) called with short-read mapping^8,19–22^, or use benchmark genomes that have curated, phased variation call sets^23–26^. Both methods rely on a reference sequence as the primary source to detect heterozygous variations. However, these reference sequences are incomplete, and recent studies have demonstrated the shortcomings of the current human reference genome and variant call sets^8,20,27^. For example, the highly variable major histocompatibility complex (MHC) is excluded from the Genome in a Bottle (GIAB) and Global Alliance for Genomic Health (GA4GH) reference panels^25^ due to its repetitive nature and need of a specialized mapping strategy to account for the high allelic diversity^28^. Moreover, reference-guided strategies require significant manual curation and effort and will not scale as large cohort sequencing projects become common^29,30^. This validation strategy is also not applicable to any species without a curated and complete reference^31^.

To overcome these limitations, we developed Merqury, which generates assembly assessment metrics using k-mers alone. Merqury compares a set of k-mers derived from unassembled, high accuracy sequencing reads to a genome assembly for evaluation. The generated assembly metrics include consensus quality (QV), k-mer completeness, and visual aids for representing the copy number spectrum and k-mer coverage across the assembly. When parental genomic sequences are available (either assembled or unassembled), Merqury can also report haplotype completeness, phase block statistics (including switch error rates), and visual representations of phase consistency for the child’s genome. We show that k-mer-based assembly validation produces comparable or better results than existing methods, such as BUSCO gene completeness and mapping-based measurements.

## Results

To demonstrate the ability of Merqury to evaluate the accuracy, completeness, and phasing of an assembly, we first applied it to an *Arabidopsis thaliana* F1 hybrid^20^, for which the parental strains (Col-0 and Cvi-0, simplified as Col and Cvi) have also been sequenced. For a comparison of multiple assemblies, we demonstrate Merqury on haplotype-resolved (TrioCanu^10^), pseudo-haplotype (FALCON-Unzip^20^), and mixed-haplotype (Canu^32^) assemblies of this hybrid genome. Total assembly size is typically used as a rough measure of haplotype completeness. For example, the TrioCanu haplotype assemblies have similar total bases, 122∼124 Mbp (**Table 1**), close to the expected haploid genome size of 130 Mbp, indicating the haplotype assemblies are well balanced (assuming haplotype specific bases are evenly inherited). In comparison, the primary assembly of FALCON-Unzip has ∼35 Mbp bases more than the alternate assembly. However, it is difficult to understand where this difference originates from the assembly size alone. The mixed-haplotype Canu assembly, in comparison, is ∼100 Mbp larger than the expected genome size of 130 Mb. Again, we can assume this assembly resolved both haplotypes, but since the haplotypes have been combined in a single assembly we cannot know the composition from the size and continuity measures alone. In the following sections we describe how Merqury’s statistics and plots can be used to dissect and understand these assemblies.

**Table 1.** Merqury quality and completeness statistics for example *Arabidopsis thaliana* and human genome assemblies.

|  | Assembly | Continuity |  |  |  | QV |  |  | k-mer completeness |  |  |  | BUSCO (%) |  |  |  |
| --- | --- | --- | --- | --- | --- | --- | --- | --- | --- | --- | --- | --- | --- | --- | --- | --- |
|  |  | Total | # Contigs | Max | NG50 | **Meryl** | **Mash** | Map | **All** | **Hap1** | **Hap2** | **PPV** | Comp. | Dup. | Frag. | Mis. |
| ***A. thaliana***<br>**Col x Cvi**<br>**F1** | **TrioCanu** |  |  |  |  |  |  |  |  |  |  |  |  |  |  |  |
|  | Col | 124 | 215 | 13.1 | 4.6 | 35.1 | 34.6 | 37.5 | 83.8 | 99.5 | 0.7 | 99.3 | 98.2 | 1.6 | 0.5 | 1.3 |
|  | Cvi | 122 | 163 | 12.2 | 5.6 | 36.4 | 35.9 | 38.6 | 83.6 | 1.1 | 99.4 | 98.9 | 98.0 | 1.4 | 0.4 | 1.6 |
|  | Col + Cvi | 246 | 378 | 13.1 | 5.5 | 35.7 | 33.0 | 38.0 | 98.3 | 99.7 | 99.4 | 99.1 | 98.2 | 78.8 | 0.2 | 1.6 |
|  | **FALCON-Unzip** |  |  |  |  |  |  |  |  |  |  |  |  |  |  |  |
|  | pri | 140 | 172 | 13.3 | 8.0 | 34.8 | 33.9 | 36.9 | 87.1 | 65.2 | 59.8 | N/A | 98.1 | 6.2 | 0.3 | 1.6 |
|  | alt | 105 | 248 | 11.6 | 4.3 | 38.3 | 37.9 | 39.9 | 74.5 | 38.2 | 40.6 | N/A | 93.1 | 2.0 | 0.3 | 6.6 |
|  | pri + alt | 245 | 420 | 13.3 | 6.9 | 36.0 | 33.5 | 37.9 | 97.8 | 99.1 | 98.2 | N/A | 98.1 | 93.2 | 0.3 | 1.6 |
|  | **Canu** | 248 | 2,368 | 6.9 | 2.3 | 29.3 | 26.8 | 27.2 | 95.7 | 90.5 | 90.8 | N/A | 97.6 | 61.5 | 0.4 | 2.0 |
| ***H. sapiens***<br>**NA12878** | **TrioCanu** |  |  |  |  |  |  |  |  |  |  |  |  |  |  |  |
|  | mat | 2,749 | 7,388 | 9.0 | 1.1 | 31.3 | 30.4 | 34.1 | 94.0 | 90.7 | 0.9 | 99.1 | 86.5 | 0.8 | 6.9 | 6.6 |
|  | pat | 2,743 | 7,252 | 11.5 | 1.1 | 31.0 | 30.1 | 34.1 | 93.7 | 1.0 | 91.0 | 98.8 | 85.1 | 0.7 | 7.8 | 7.1 |
|  | mat + pat | 5,492 | 14,640 | 11.5 | 1.1 | 31.1 | 27.5 | 34.1 | 98.2 | 90.9 | 91.2 | 99.0 | 90.0 | 47.3 | 5.0 | 5.0 |

All bases in the continuity stats are in Mbp. Haploid genome size of 130 Mbp and 3.2 Gbp was used for NG50 in the *A. thaliana* and *H. sapiens* NA12878 haploid assemblies, respectively, with twice the haploid genome size for combined assemblies. Mercury-specific column headers are in bold, including k-mer-based quality (QV) and completeness estimates. Mercury includes both exact (Meryl) and approximate (Mash) methods for measuring k-mer QV, while completeness uses only the exact k-mer counting method. Consensus QV scores are Phred-scaled where $QV = -10 \log_{10} E$ for a probability of error $E$ at each base in the assembly. K-mer completeness is measured by the fraction of all distinct, reliable k-mers (all) and haplotype specific k-mers. Hap1 and Hap2 corresponds to Col and Cvi in *A. thaliana*, maternal and paternal in NA12878. BUSCO v3 was run with embryophyta\_odb9 (n=1440) for *A. thaliana* and mammalia\_odb9 (n=4104) for NA12878. Metrics generated with Mercury are highlighted in bold. Map. Mapping; Comp., Complete; Dup. Duplicated; Frag. Fragmented; Mis. Missing.

### Copy number spectrum

We counted k-mers from Illumina whole-genome sequencing of the *A. thaliana* F1 hybrid, as well as from each assembly, using Meryl, a k-mer counting tool we extended to support k-mer set operations for Merqury (**Methods**). The copy number spectrum plot, known as “spectra-cn” plot^17^ tracks the multiplicity of each k-mer found in the Illumina read set (**Fig. 1a**) and colors it by the number of times it is found in a given assembly (**Fig. 1b**). The result is a set of histograms relating k-mer counts in the read set to their associated counts in the assembly. Here the Illumina dataset (which we will refer to as the “read set”) was sequenced to an average coverage of 45x, so we expect a histogram peak near x=45 corresponding to k-mers present in both haplotypes, and a peak at half coverage (x=22) representing k-mers found on only one haplotype (**Fig. 1a**, in practice, these peaks are shifted slightly lower due to the effects of sequencing error and sampling only *l*-*k*+1 *k*-mers for each *l*-sized read). We refer to these as 2-copy and 1-copy k-mers, respectively, to indicate the number of times they appear in the true genome. Additional peaks may appear for polyploid genomes, but for the remainder of this paper we will assume a diploid genome.

**Fig. 1.**
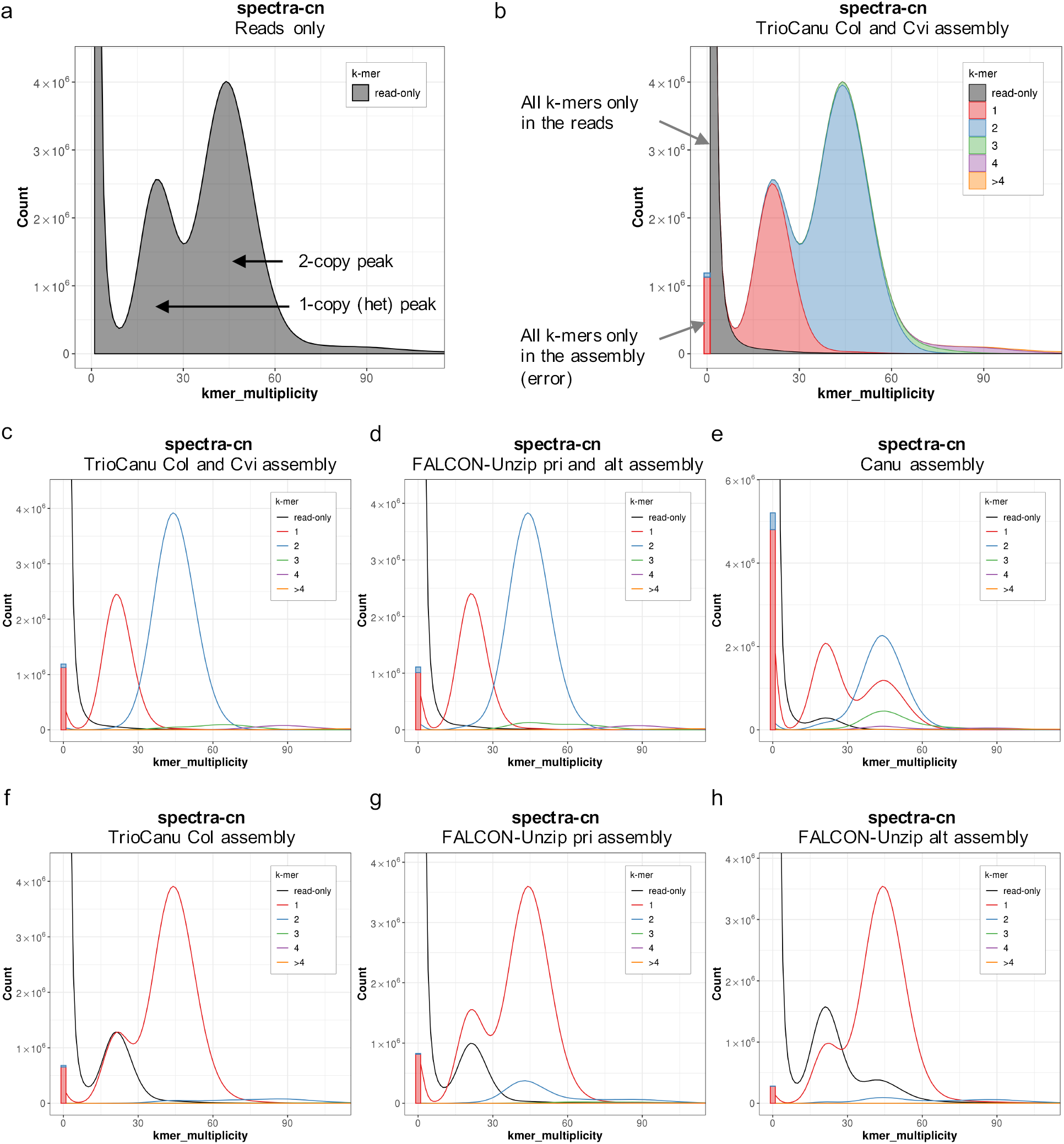
Merqury copy number spectrum plots for haploid and diploid assemblies of an *Arabidopsis thaliana* F1 hybrid genome. (a) Histogram of k-mer multiplicity collected from Illumina reads. By default, Merqury connects the midpoint of each histogram bin with a line, giving the illusion of a smooth curve. The first peak represents 1-copy (heterozygous) k-mers in the genome, and the second peak represents 2-copy k-mers originating from homozygous sequence or haplotype-specific duplications. Depth of sequencing coverage determines where these peaks appear. In this example, sequencing coverage is approximately 45x, corresponding to the 2-copy peak. (b) Copy number spectrum (spectra-cn) of the same k-mers in **a** plotted as stacked histograms colored by the copy numbers found in the combined TrioCanu assembly. The assembly k-mers absent from the read set (likely to be base errors in the assembly) are plotted as a bar at zero multiplicity, colored by the copy numbers found in the assembly. (c) Unstacked histograms of **b** for visualizing the distribution of k-mer counts per copy numbers found in the assembly. This plot shows an ideal pseudo haplotype assembly. (d) Spectra-cn plot of the combined FALCON-Unzip assembly. (e) Spectra-cn plot of the mixed-haplotype Canu assembly. Missing single copy k-mers (black) and k-mers from artificial duplications (green, purple, yellow in 30–60x) are noticeable. Note this assembly was not polished and so has a larger error k-mer bar. (f) Spectra-cn plot of the TrioCanu Col haplotype assembly. Half the single copy k-mers are missing and found in the other haplotype (black). Two-copy k-mers are found once (red) in each haplotype assembly. (g) Spectra-cn plot of the FALCON-Unzip primary assembly. (h) Spectra-cn plot of the FALCON-Unzip alternate assembly.

Thus, when a k-mer is found approximately 22 times in the *A. thaliana* read set, we expect it to be found only once in the assembly, as it is likely a 1-copy, haplotype-specific (heterozygous) sequence (**Fig. 1b**). In the spectra-cn plot, k-mers are colored based on their count in the assembly. For a complete haplotype-resolved assembly, where even the homozygous part of the genome is included in both haplotypes, we expect most k-mers in the 2-copy peak to be found twice in the assembly (**Fig. 1b**). For partially phased assemblies, 2-copy k-mers may be found either once or twice in the assembly (e.g. **Fig. 2**), depending on which homozygous sequences of the genome were separated and which were collapsed. In contrast, a pseudo-haplotype collapses homozygous alleles, so 2-copy k-mers are expected to appear only once in the assembly. One notable exception is haplotype-specific duplications, which can occur in two copies on the same haplotype, and thus may also appear in two copies in a pseudo-haplotype assembly.

**Fig. 2.**
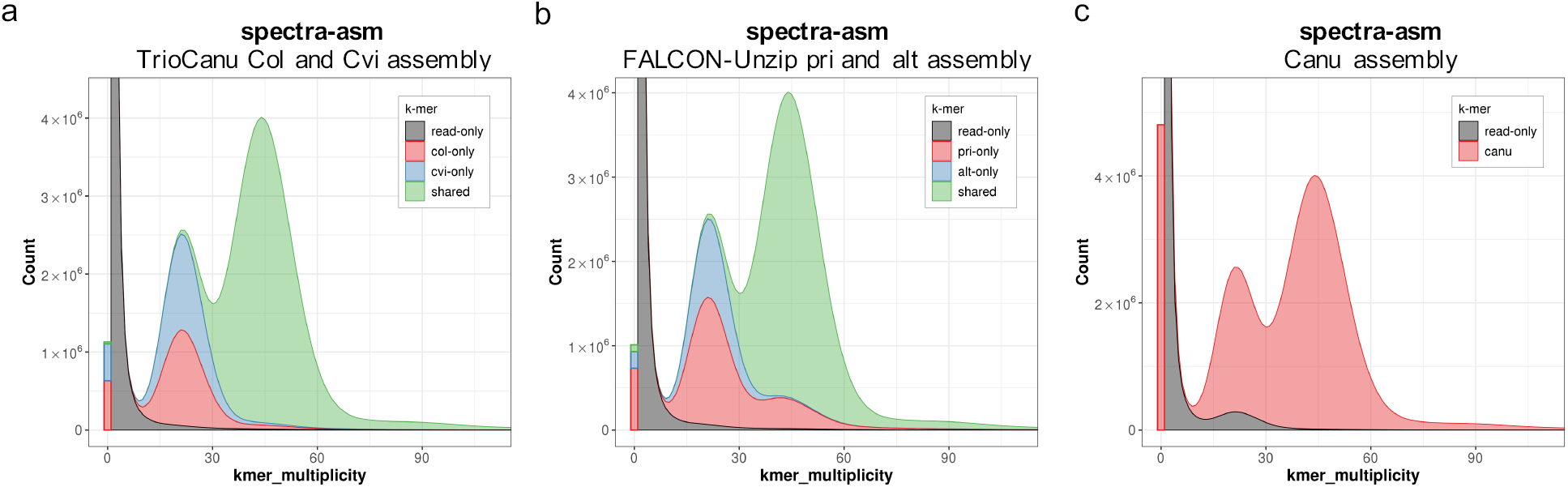
Merqury assembly spectrum plots for evaluating k-mer completeness. K-mers are colored by their presence in the reads and primary/alternate assemblies. (a) Distinct k-mer assembly spectrum (spectra-asm) plot of both TrioCanu Col and Cvi haplotype assemblies. This plot shows the assembly specific (red and blue) and shared portion of k-mers (green). (b) Spectra-asm plot of the FALCON-Unzip assembly. The primary assembly has more k-mers (red) compared to the alternate assembly (blue). (c) Spectra-asm plot of the Canu assembly which is a mixture of both haplotypes. A small fraction of 1-copy k-mers is found only in the reads (black peak around 12∼30x), which represents heterozygous variants missing from the assembly.

Assuming no serious biases in the read set, a clean spectra-cn plot is a necessary, but not sufficient, condition of assembly quality. Quoting KAT author Bernardo Clavijo “if your spectra look right, your assemblies could still be wrong, but if your spectra look wrong your assemblies can’t be right.”. That is to say, certain structural errors may not be reflected by the spectra, but any conflict with the copy numbers of the read set indicate assembly problems. The k-mers found only in the read set (black) at low frequency are almost always indicative of sequencing error in the read set, however higher-frequency k-mers found only in the read set indicate missing sequences in the assembly (e.g. black k-mers within the 1- or 2-copy peaks). Likewise, any k-mers with a higher copy number in the assembly than would be predicted by the read set are indicative of artificial duplications in the assembly, e.g. see the 2-copy k-mers appearing three times in the Canu assembly shown in **Fig. 1e**.

The bar at the origin of the plots represents k-mers found only in the assembly. From these k-mers, we can estimate an assembly consensus quality value (QV), which represents a log-scaled probability of error for the consensus base calls (**Methods**). Higher QVs indicate a more accurate consensus, where Q30 corresponds to 99.9% accuracy, Q40 to 99.99%, etc. The trio-binned assembly has QV scores of 35∼36 for each haploid assembly, and 35.7 for the combined version. The FALCON-Unzip assembly has a similar QV score of 35∼38 for each haplotype, and 36 for the combined version. The error k-mer bar for Canu is much higher than the other two diploid assemblies, as we omitted signal-level polishing^33^ from this assembly to show an intermediate assembly product. The estimated QV for this assembly is 29.3 (**Table 1**).

To better visualize the k-mer distribution by each copy number found in an assembly, we also provide unstacked versions of the spectra-cn histograms (**Fig. 1c-e**). The 1-copy (heterozygous) k-mers appear once in the combined TrioCanu assemblies (red), and 2-copy k-mers twice (blue) as expected (**Fig. 1c**). The partially phased FALCON-Unzip assembly shows a similar distribution to TrioCanu (**Fig. 1d**), indicative of good k-mer completeness. The lower fraction of 2-copy k-mers found three times in the assembly (green hump under the 2-copy peak) indicates fewer false duplications in the TrioCanu compared to FALCON-Unzip and Canu.

When generating the same spectrum on the mixed-haplotype Canu assembly (**Fig. 1e**), we can see the assembly has only partially assembled both haplotypes and there is a higher fraction of k-mers (37 kbp) missing from the assembly (black). In addition, the assembly has artificial duplications inflating the assembly size. The plot also shows fewer 2-copy k-mers (blue peak) compared to the other assemblies, and a significant fraction of 2-copy k-mers appearing only once in the assembly (second red peak), suggesting the mixed-haplotype assembly has partially collapsed the haplotypes. Based on the number of 2-copy k-mers found twice in the assembly, we estimate 42.7 Mbp of homozygous sequence remains un-collapsed, typically at the boundary of heterozygous and homozygous alleles. This partial separation is also evident in the higher number of k-mers appearing three (green), four (purple), and more (yellow) times in the assembly, representing 10.8 million artificially duplicated k-mers that would need to be identified and removed to form a fully collapsed, haploid assembly. This example highlights the benefits of the TrioCanu and FALCON-Unzip approaches for this heterozygous genome. Additional processing with a tool such as purge_dups^34^ would be required to convert Canu’s mixed haplotypes into a pseudo-haplotype.

The spectra-cn plots can also be useful for evaluating haploid assemblies, e.g. of a single haplotype from a diploid genome. When plotting the same k-mer spectrum on one haplotype (“Col” in this case), we can see both 1-copy and 2-copy k-mers are now observed just once in the assembly (**Fig. 1f**, red histogram). This is because the read set represents the full diploid genome, while the assembly isolates a single haplotype. The usual 1-copy peak is now exactly half the size and perfectly overlaps with a peak of missing k-mers (black) that belong to the other haplotype (“Cvi”). In comparison, the pseudo-haplotype FALCON-Unzip primary and alternate assemblies (**Fig. 1g-h**) show imbalanced peaks, with more assembly k-mers appearing in both the 1- and 2-copy peaks than expected. This suggests that the FALCON-Unzip is erroneously including sequences from both haplotypes into the primary pseudo-haplotype. A similar portion of 1-copy k-mers are missing from the alternate assembly (imbalanced red and black peaks), suggesting that the alternate haplotype is missing some heterozygous variants.

### Assembly spectrum

In the above FALCON-Unzip pseudo-haplotype example (**Fig. 1g-h**), it is possible to infer that the missing sequences in the alternate assembly are likely found in the primary assembly. However, if there are shared sequences between the two assemblies, it is difficult to know the exact sequence composition. To better address this question, we introduce a new method to show the shared and specific k-mers in each assembly (spectra-asm), instead of showing the overall copy-numbers (**Fig. 2**). This plot is helpful for measuring diploid assembly completeness as it shows the fraction of k-mers specific to both the primary and alternate assemblies. For example, a perfectly assembled diploid genome is expected to have a balance of k-mers specific to each haplotype representing the heterozygous alleles (exceptions to this include sex chromosomes of different size). The spectra-asm of the TrioCanu combined assembly (**Fig. 2a**) shows such an example, where 1-copy k-mers are specific to each haplotype assembly (red and blue), and the 2-copy k-mers are shared by both assemblies (green). In comparison, the FALCON-Unzip assembly is imbalanced, with more 1-copy and 2-copy k-mers in the primary assembly than expected (**Fig. 2b**). This imbalance is also evident from Merqury’s k-mer completeness metrics for the primary and alternate assemblies (**Table 1**, completeness “all”). By stacking the histograms, we can confirm the primary assembly contains all the missing k-mers from **Fig. 1h** in the 2-copy peak. Compared to TrioCanu and FALCON-Unzip, Canu does not partition its output into primary and alternates, and so k-mers from both haplotypes are present in the combined assembly. However, the spectra-asm plot shows a few 1-copy k-mers are missing from the assembly (**Fig. 2c**). This is in concordance with the lower k-mer completeness score in the Canu assembly (95.7%) compared to the other diploid assemblies (98.3% and 97.8%, **Table 1**), and indicates that some heterozygous variants are missing, perhaps as a result of the low-quality, unpolished consensus.

### Haplotype-specific k-mers (hap-mers) from trios

We define a “hap-mer” as a haplotype-specific k-mer that appears exclusively, one or more times, on a single haplotype of the genome. When parental genomes are available, we can use inheritance to estimate a set of hap-mers for the child and evaluate haplotype completeness of the assembly. Merqury identifies hap-mers as the set of inherited, parental-specific k-mers (**Fig. 3a**). Using the parental-specific markers alone may be sufficient for many cases; however, we have found it useful to specifically consider the inherited markers, as only half of the parental specific k-mers are inherited and the non-inherited markers may match an erroneous k-mer in the assembly by chance. Note, this is still only an estimate of the true set of hap-mers in the child’s genome, which can also be affected by *de novo* variants or heterozygous variants that were differentially inherited from the parents. We have implemented efficient k-mer set operations (union, intersection, subtraction, etc.) within Meryl for computing hap-mers and other useful k-mer sets (**Fig. 3b, Methods**). For the *A. thaliana* F1 hybrid genome, we identified hap-mers directly from the genomes of the parental strains. K-mers were grouped based on their presence in the F1 reads alone, the maternal haplotype, the paternal haplotype, or both (**Fig. 3c**). Because of the high heterozygosity of the F1 (estimated at 0.99% by GenomeScope^35^), many of the F1’s k-mers are hap-mers. A human genome, in comparison, has many fewer hap-mers, with most of the k-mers shared between both haplotypes (**Fig. 3d**).

**Fig. 3.**
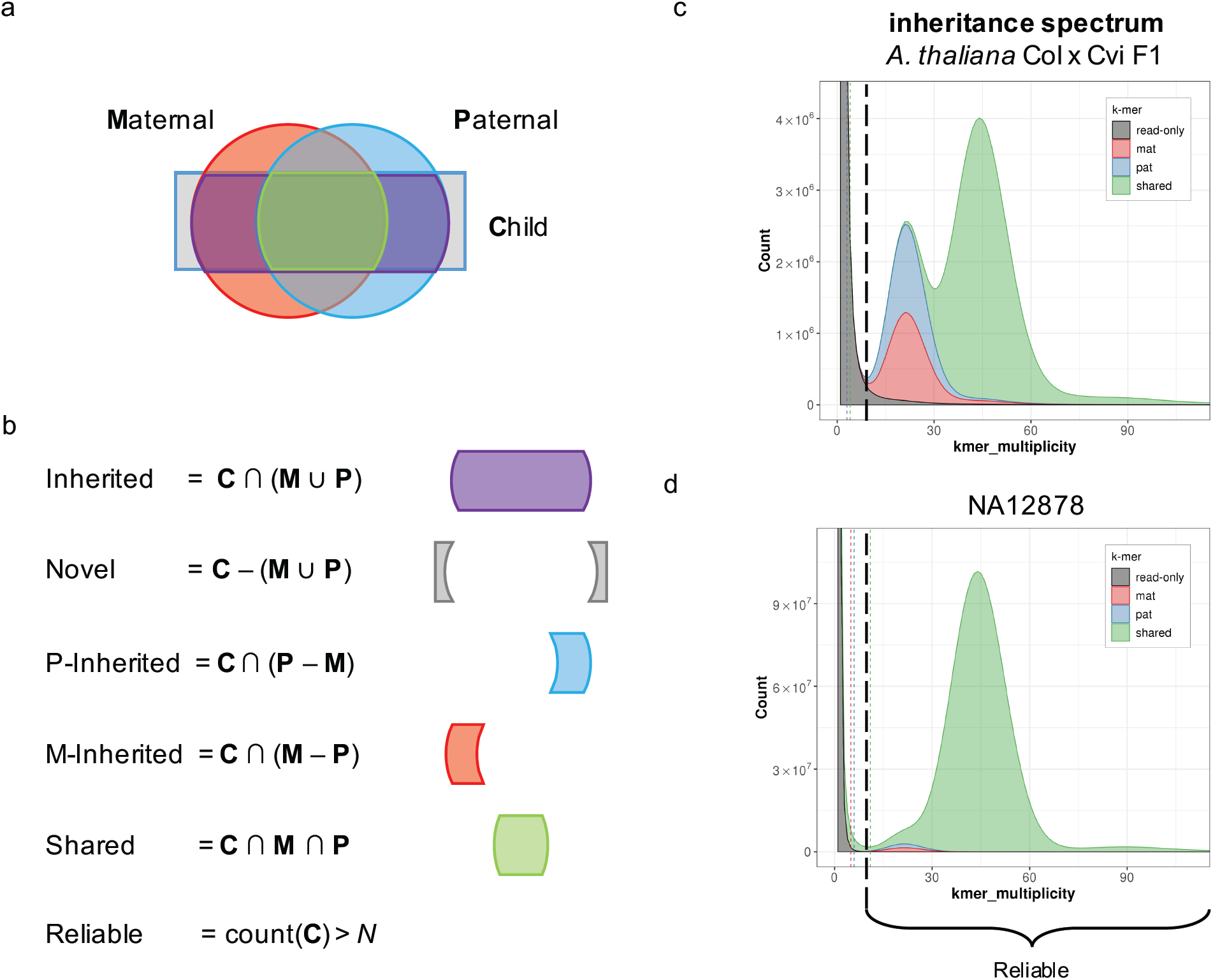
Merqury set operations for generating haplotype specific hap-mers and reliable k-mers. (a) Venn diagram of maternal, paternal, and child k-mer sets. Inherited haplotype-specific k-mers (hap-mers) are estimated from the parental and child k-mer sets. Roughly half of the parental-specific k-mers are inherited by a child. (b) Set operation examples used in Meryl to compute hap-mers and other k-mer sets. (c-d) Stacked k-mer multiplicity of the child’s read set, colored by inheritance. K-mers are colored by maternal (red) paternal (blue) and shared between parents (green). K-mers only seen in the child’s reads (black) are mostly from low-copy sequencing errors or k-mers arising from *de novo* variants in the child. Reliable k-mer thresholds used for generating completeness scores for all k-mers (black) and hap-mers (red and blue) are marked by dashed lines.

### Evaluating phasing completeness with hap-mers

Hap-mers are used to determine phase blocks in Merqury, where a block is defined to be a consistent set of markers originating from the same haplotype. To account for minor base-level errors in the assembly, we do allow some short-range switches to occur within a block, so long as the phase switches back shortly thereafter (**Fig. 4a**). A benefit of this k-mer approach is that Merqury does not need to rely on the phase blocks as identified by the assembler or a reference variant callset, and can quickly compute the blocks on each assembly directly using only the observed haplotype markers. Applying this method to the TrioCanu assembly reported NG50 phase block sizes of 3.6 Mbp and 5.5 Mbp with 0.3% per-block switch rate when allowing at most 100 consecutive switches within 20 kbp (**Fig. 4b**). The FALCON-Unzip assembly has slightly shorter phase block sizes of 3.1 Mbp and 2.5 Mbp with a similar switch error rate of 0.3%. The Canu assembly had more frequent long-range switches among haplotypes, resulting in NG50 phase blocks of 100 kbp.

**Fig. 4.**
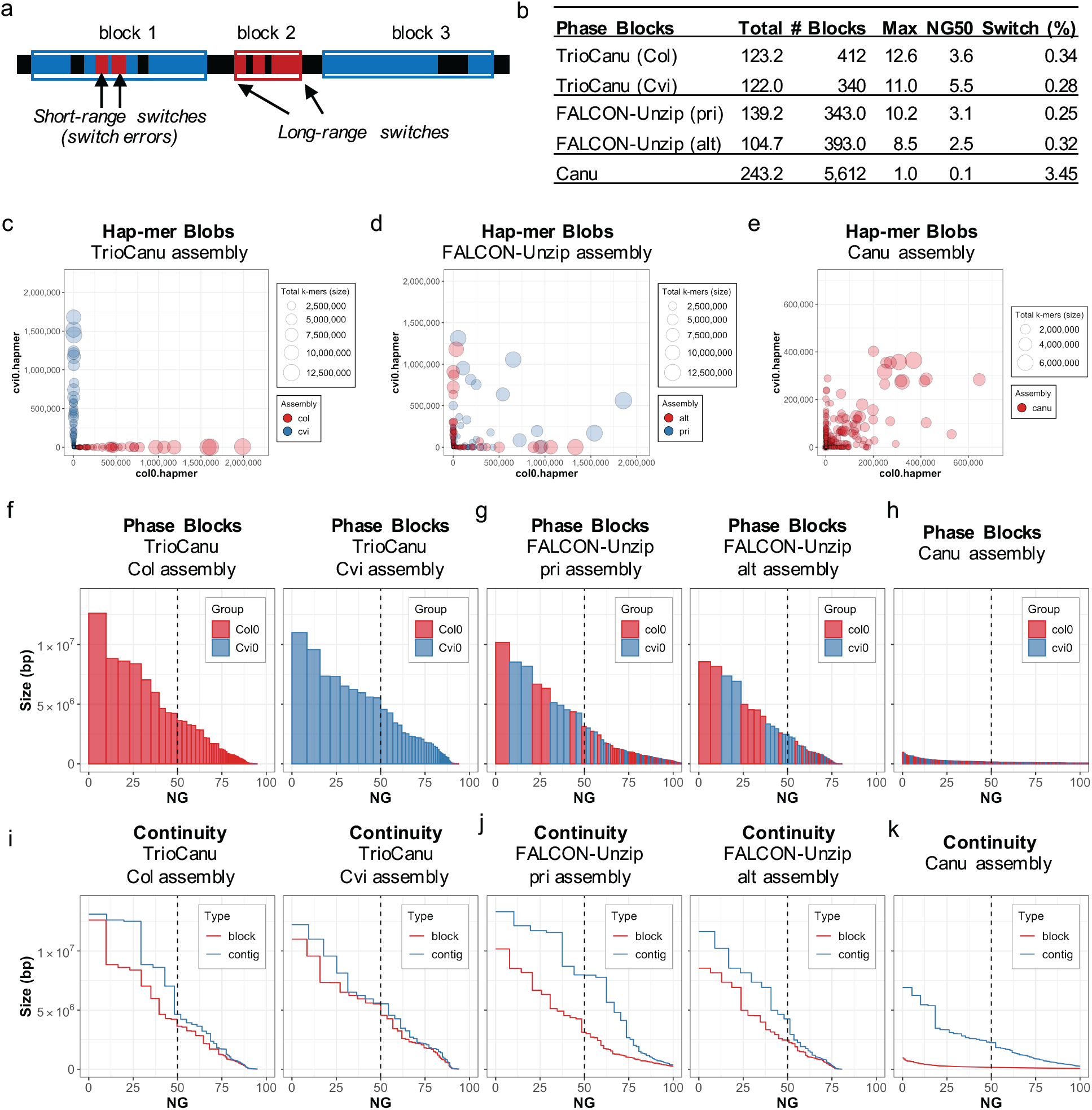
Merqury hap-mer plots for evaluating haplotype phasing. (a) Example of phase blocks and switches. Blue and red bars are paternal or maternal hap-mers found in the assembly. A phase block is defined by at least two hap-mers (markers) from the same haplotype. Short-range switches are allowed in between markers, in defined ranges. Two consecutive red markers within a certain range are marked as short-range switches and counted for switch errors in block 1. As the red markers are consecutively found over a certain range, or in numbers above a certain threshold, a separate block is formed. Each switch between blocks is counted as a long-range switch. (b) Phase block statistics of the haploid assemblies with switch errors, allowing at most 100 switches within 20 kbp. (c) Hap-mer blob plot of the TrioCanu assembly. Red blobs represent Col haplotype contigs, while blue blobs are the Cvi haplotype. Blob size is proportional to contig size, and each blob/contig is plotted according to the number of contained Col (x values) and Cvi (y value) hap-mers. Col-specific k-mers are found in the Col assembly with almost no Cvi-specific k-mers, while Cvi k-mers are found in the Cvi assembly with almost no Col k-mers. (d-e) Blob plots for FALCON-Unzip and Canu assemblies show that most contigs are a mix of sequences from both haplotypes, but FALCON-Unzip preserves phase within its alternate contigs, as designed. (f) Phase block NG* plots of the haplotype resolved Col (left) and Cvi (right) assembly, sorted by size. X-axis is the percentage of the genome size (*) covered by phase blocks of this size or larger (Y-axis). Blocks from the wrong haplotype are very small and almost entirely absent. (g-h) Phase block NG* plot of the (g) FALCON-Unzip and (h) Canu assemblies. Col and Cvi phase blocks are distributed evenly, as is typical for pseudo-haplotype assemblies. (i-k) Phase block and contig NG* plots show the relative continuity of (i) TrioCanu, (j) FALCON-Unzip, and (k) Canu assemblies. Phase block sizes are similar to the contig sizes in **i**. Phase blocks are much shorter than the contigs in **k**, because of the frequent haplotype switches in the contigs.

Visualizing hap-mer presence in each haplotype assembly is also useful to detect overall phase consistency. When counting Col- and Cvi-specific k-mers in contigs of the TrioCanu assembly (**Fig. 4c**), each contig was successfully separated by haplotype as expected. That is, the Col markers were observed in the Col haplotype assembly, with almost no contaminating Cvi markers, and vise versa. As such, a haplotype resolved assembly with almost no haplotype switches is expected to have a similar plot with each blob (contig) close to the corresponding plot axis. The FALCON-Unzip alternate contigs maintain haplotype consistency (**Fig. 4d**), but the primary pseudo-haplotype contigs are a mixture of both haplotypes. The Canu assembly appears to mix haplotypes in all but the smallest contigs (**Fig. 4e**), and does not partition the resulting assembly into primary and alternate contig sets with post-processing with Purge_dups.

When plotting phase blocks sorted by size, the blocks originating from the wrong haplotype were very small and almost negligible in the TrioCanu assembly (**Fig. 4f**). In contrast, the phase blocks were highly mixed in the pseudo-haplotype assemblies, with the larger contigs being more likely to contain markers from both haplotypes (**Fig. 4g-h**). Plotting the contig and block sizes together shows that the trio-binned phase blocks are very similar in size to the trio-binned contigs (**Fig. 4i**). In comparison, the phase blocks were shorter than the contigs in the FALCON-Unzip assemblies (**Fig. 4j**), showing relatively good phasing performance. The phase blocks were much shorter in the Canu contigs, indicating frequent block switches between haplotypes (**Fig. 4k**) since Canu does not attempt to preserve long-range phasing.

Another useful feature in Merqury is that all hap-mers, erroneous k-mers, and phase blocks can be visualized as genome tracks along the assembly. **Fig. 5** shows an example of a 60 kbp region in the mixed-haplotype Canu assembly, where haplotypes are observed switching from Cvi (blue) to Col (red), resulting in numerous base errors (grey). This illustrates how failure to separate haplotypes can result in increased base error, as the two haplotypes are improperly combined into a single consensus sequence resulting in artifactual variants. Two phase blocks (Cvi and Col) were found in this region using the default threshold of 100 consecutive k-mer switches allowed per every 20 kbp. However, phase blocks can be more stringently measured by defining the short-range switch allowance (e.g. 10 per 20 kbp, **Fig. 5**, bottom track), resulting in lower NG50 phase block size (100 kbp decreases to 33 kbp). In contrast, the per-block switch error rate decreased from 3.4% to 0.47%, making each block a more reliable haplotype. Note the per-block switch error rate is defined as the fraction of k-mer markers within a block that are assigned to the wrong haplotype, thus accounting for all short-range marker switches in a block.

**Fig. 5.**
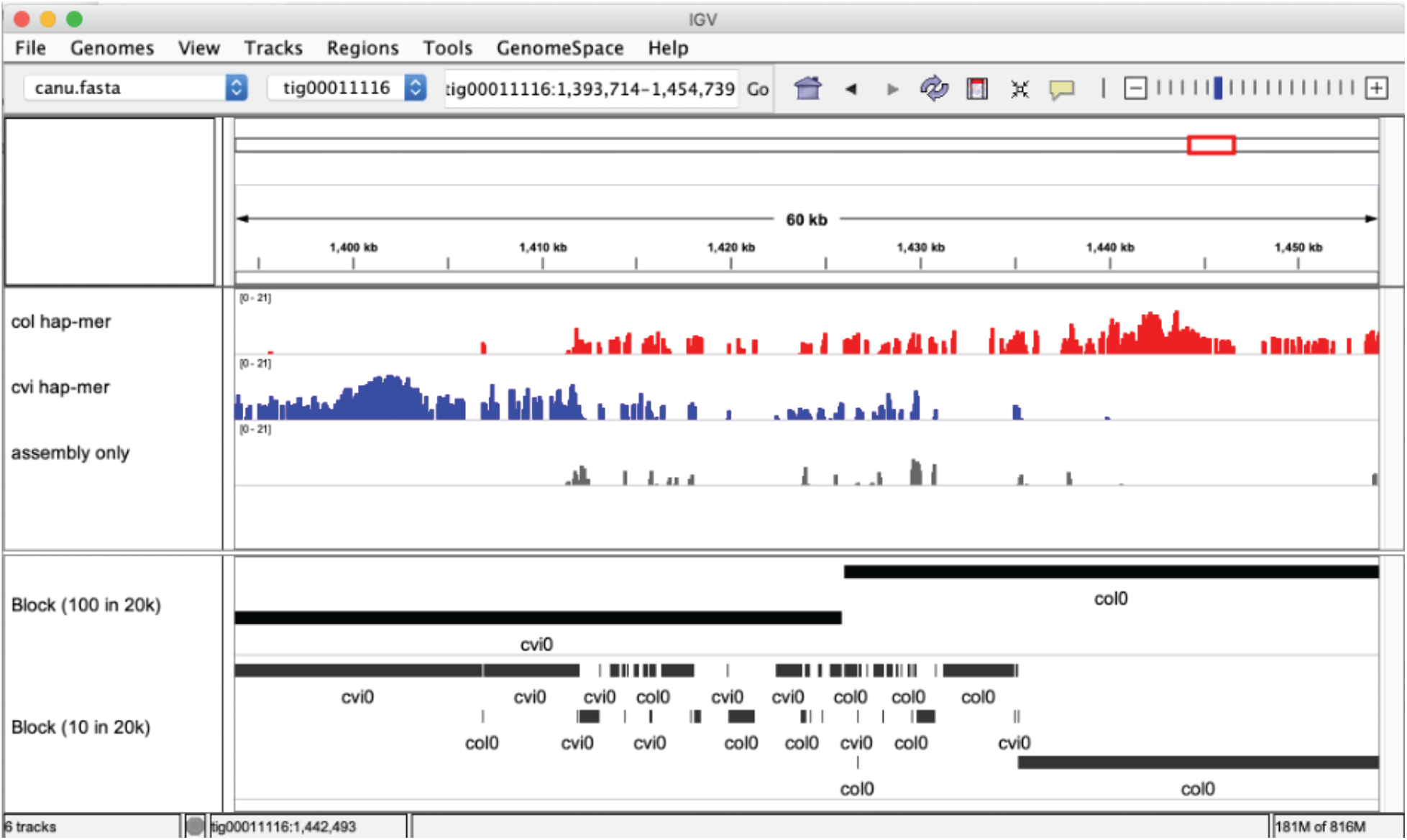
Example k-mer and phasing tracks provided by Merqury. Hap-mer density is provided in tiled data format (.tdf) browsable with the Integrated Genomics Viewer (IGV)^36,37^. This figure shows a region where haplotype blocks are switching within one of the Canu contigs. Hap-mer tracks show haplotype switches from Cvi (blue) to Col (red), along with k-mers found only in the assembly (grey), which are likely caused by erroneous consensus bases. Phase blocks (black) are shown for both relaxed (100 consecutive switches allowed per 20 kbp) and strict (10 per 20 kbp) switching thresholds.

### Benchmarking on a human genome (NA12878)

To benchmark Merqury on a large genome, we applied it to the TrioCanu human (NA12878) assembly from Koren *et al*.^10^ and estimated the consensus quality as Q31 for each haplotype (**Table 1**). The alternative variant calling approach reported 2.1 million bases of errors, resulting in a QV estimate of 34.1. However, the mapping-based approach excluded 212.2 Mbp of assembled sequence because of too few (<3x) or too many (>600x) aligned reads. We argue that Merqury’s k-mer based approach is both more efficient and more accurate for evaluating consensus accuracy.

Merqury required only 14.9 CPU h (9.1 h for k-mer counting, 4.7 h for merging, 1.1 h for statistics) to evaluate QVs for both haplotypes. In contrast, the mapping-based QV estimate took 338.3 CPU h (2.2 h for indexing, 308 h for mapping, 12.6 h for merging, 12.8 h for variant calling, and 2.5 h for coverage calculation and QV estimates). By excluding low and high-coverage regions of the assembly, the mapping-based approach ignores regions of the assembly likely to be enriched for error. For example, low coverage regions can be caused by regions of high error rates that makes it difficult to map short reads. High coverage regions are typically caused by repeats that can be collapsed, and therefore incorrect, in the assembly. Thus, a substantial number of errors may be excluded from the accuracy statistics if one considers only the mappable portion of the assembly. This matches with our observation that the mapping-based estimates always overestimate QV compared to the k-mer based approach (**Table 1**).

Exact k-mer counting is currently the most resource intensive step of Merqury, requiring a maximum 21 GB of memory using 25.5 GB of disk space on NA12878 (**Table 2**). While this step can be parallelized across multiple nodes and cores, QV statistics can be also estimated from subsampled k-mers with lower memory and disk requirements using Mash Screen^38^. Because Merqury’s QV estimation is based on Mash’s k-mer containment score (**Methods**), the Meryl and Mash counting methods are interchangeable. In comparison to Meryl, Mash streams sequencing reads from disk and compares them against only a small subset of k-mers in the assembly. This avoids the need for a large table of k-mers, but at the same time ignores copy number information. As a result, we observed that Mash QV estimates were slightly lower than exact counting (Meryl) for each haplotype, and even lower when both haplotypes were combined (**Table 1**). This is because the shared k-mers between the two haplotype assemblies are considered only once by Mash, resulting in an underestimate of the QV score (e.g. if a 2-copy k-mer appears in just one haplotype, it is considered “correct” by Mash). The Mash approach also cannot investigate positional base errors (**Figure 5**) and many of the other analyses presented here, but is provided as an alternative to Meryl for QV estimation in cases where disk and memory resources are limited.

**Table 2.**
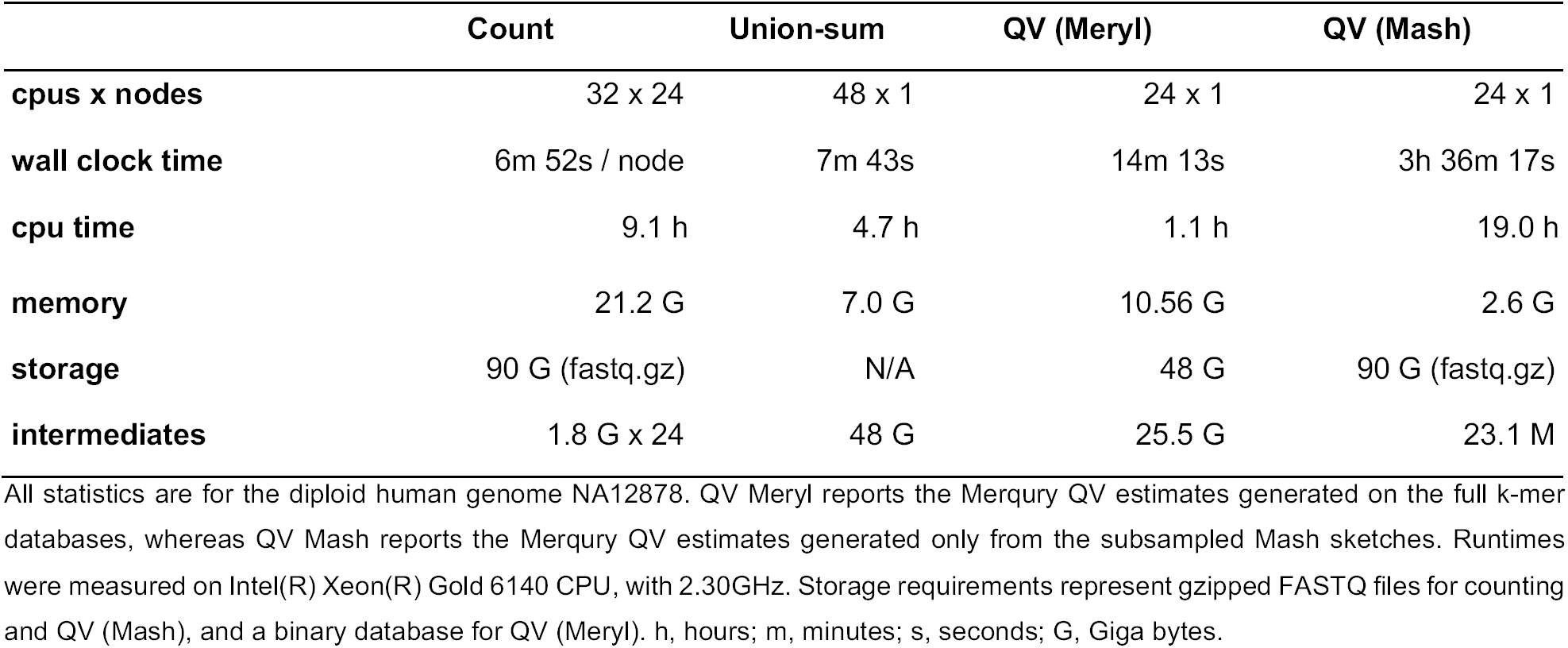
Merqury runtime, memory, and disk requirements for a human genome.

Next, we evaluated k-mers for NA12878. The hap-mer spectrum of NA12878 revealed a higher fraction of shared k-mers in the 1-copy peak (**Fig. 3d**), indicating that some heterozygous variants in the child are shared by both parents. We do not see such a strong effect in *A. thaliana*, because the parents were heavily inbred and contained few heterozygous variants of their own. In contrast, the *A. thaliana* F1 hybrid was deliberately outbred, which is evident by the dramatically taller 1-copy peak versus NA12878 (0.99% vs. 0.12% heterozygosity).

Haplotype-specific k-mers are convenient to obtain haplotype precision (PPV) and recall (completeness) statistics based on how many of the expected parental k-mers are observed in the child’s haplotype-resolved diploid assembly (**Fig. 6**). To demonstrate, we built genomic k-mer databases for NA12878 and her parents, totaling 18.4 and 19.9 million inherited hap-mers for the paternal and maternal haplotypes, respectively. When comparing to the haplotype-resolved assemblies, the maternal haplotype assembly recovered 90.7% of the maternal hap-mers (**Table 1** and **Fig. 6a**), and the paternal assembly recovered 91.0% of the paternal hap-mers (**Table 1** and **Fig. 6b**). Likewise, by considering the other haplotype’s markers as false positives (i.e. paternal hap-mers found in the maternal assembly), the precision of the maternal and paternal assemblies was 99.1% and 98.8%, respectively, with only 160∼200 markers appearing in the incorrect haplotype (**Table 1**, PPV values). This evaluation excludes all k-mers found only in the assembly (errors), which if considered false positives, would further lower the precision.

**Fig. 6.**
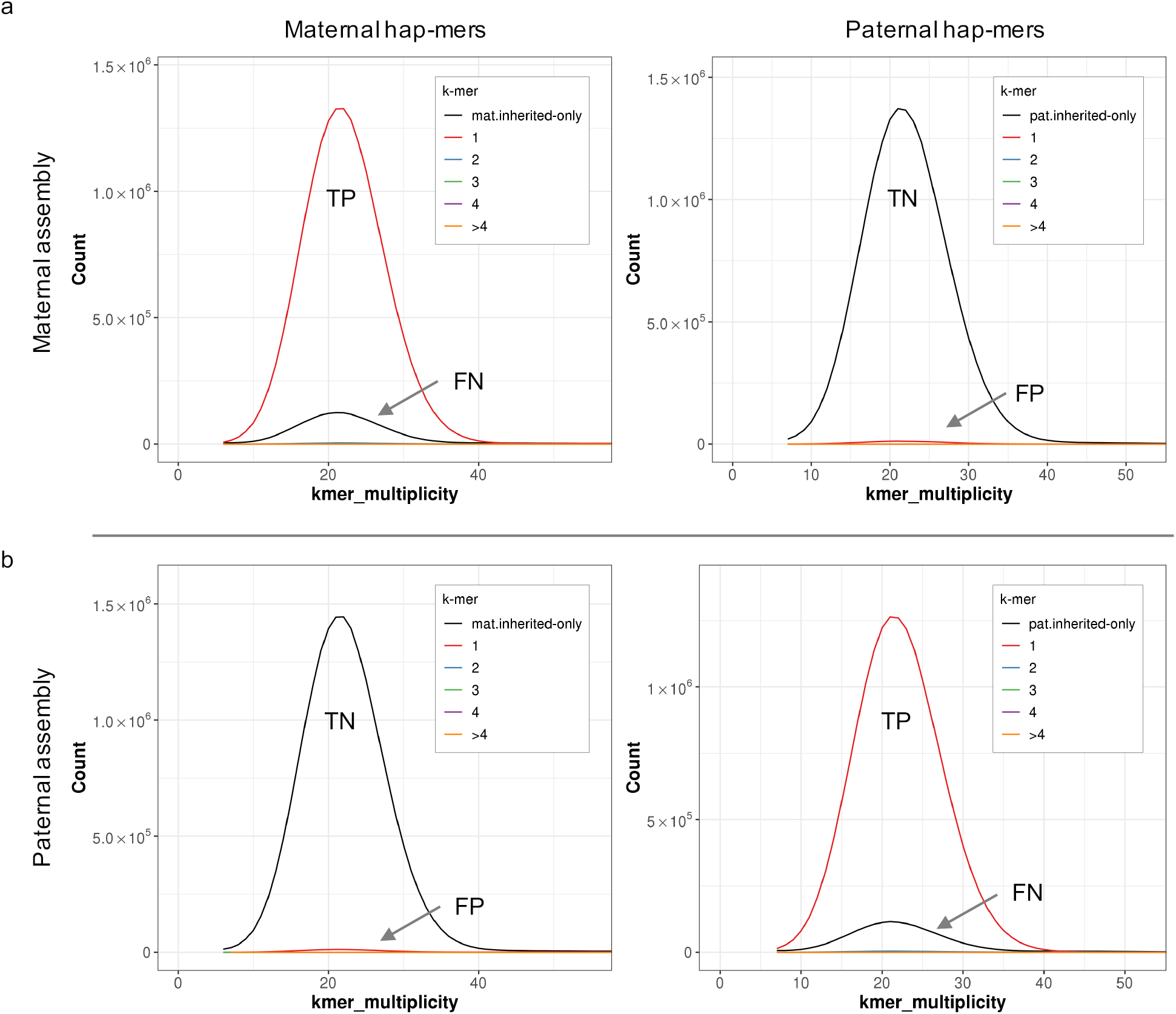
Evaluating haplotype completeness for diploid assemblies using Merqury. Spectra-cn plots of the maternal and paternal assemblies of the TrioCanu NA12878 assembly are displayed as a confusion matrix. (a) Hap-mer spectra-cn plot of the maternal assembly. All maternal hap-mers are expected to be found in the maternal (top left), with no paternal hap-mers (top right). Any maternal hap-mers not found in this assembly are missing hap-mers (top left, black). Paternal hap-mers (top right, red) are false positive phasing errors. (b) Hap-mer spectra-cn plot of the paternal assembly. This time, no maternal hap-mers should be seen (bottom left), and only paternal markers are expected to be present (bottom right). TP, true positives; FN, false negatives; TN, true negatives; FP, false positives.

To compare Merqury’s results with an alternative approach, we considered an Illumina-based platinum callset for NA12878^26^ that includes 3.4 million heterozygous SNPs in regions where both haplotype assemblies align to the reference (hg38). Calling SNPs directly from the haplotype assemblies against hg38 recovered 3.2 million variants, or 93.5% sensitivity, which is slightly higher than the Merqury based estimate of ∼91%. This discrepancy is likely due to Merqury’s ability to measure complex regions of the assembly not easily measurable by a mapping-based analysis. In contrast, the SNP-based measurement of precision was only 86.1%, likely due to the low base accuracy (Q31) of the assemblies, where consensus errors are called as false-positive heterozygous SNPs. Thus, it is important to consider both the k-mer QV and precision estimates when evaluating the accuracy of a diploid assembly.

## Conclusion

We have developed Merqury, a reference-free assembly evaluation toolkit based on efficient k-mer based methods. Merqury builds upon the k-mer spectra ideas of Mapleson *et al*^17^, and introduces novel methods and plots for measuring assembly quality (QV), completeness, and phasing. Using k-mer count spectra, Merqury can reveal copy-number errors in an assembly and accurately measure both assembly completeness and consensus quality. When parental k-mers are available, Merqury can also measure phasing accuracy and haplotype completeness. In addition to validation statistics, Merqury provides a number of graphs for interpreting assembly quality. In the process of developing Merqury, we have also extended Meryl with generalized k-mer counting, querying, and set operations that will be useful for other k-mer based analyses (**Fig. 3b**).

Merqury is able to evaluate assemblies from any sequencing technology, and works best when high-accuracy sequencing reads are available from the assembled individual and its parents. This read set serves as independent validation of the assembly, which is typically based on less-accurate, long-read sequencing. If high-accuracy reads are not available from the assembled individual, read sets from the parents can be used as a replacement for measuring quality values. This assumes all k-mers in the child are found in the parental genomes, ignoring the small fraction of k-mers from *de novo* mutations in the child. Although we currently recommend using Illumina data for the k-mer based validation, it may require special library preparation to minimize sequencing biases^39-41^. We note that Merqury’s methods are general and would be compatible with any reasonable high-accuracy, high-throughput sequencing technology.

Hap-mers are currently computed by a simple set operation, similar to trio-binning^10^. A higher portion of hap-mers are identifiable when the parents are divergent, with minimal shared ancestry. Our hap-mer spectrum plots (**Fig. 3c-d**) show the 1-copy haplotype markers that are specific to each parent, which may not be as prevalent for genomes of low heterozygosity, such as humans. Here we have used trios of diploid genomes to identify haplotype-specific markers, but Merqury’s methods are extensible to polyploid genomes and it is possible to identify hap-mers using orthogonal datasets (e.g. Hi-C, Strand-seq). To support alternative k-mer classification methods, Merqury is designed to receive any pre-computed hap-mer set as input.

We argue that Merqury’s k-mer-based method provides better haplotype completeness estimates, because it does not rely on a reference genome. Mapping to a reference can be biased by mis-mapping to repetitive or low-quality regions of the assembly. Moreover, k-mers naturally capture heterozygous insertion and deletion variants and are thus immune to the problems of calling these types of variants with a reference mapping approach. For example, consortiums such as the GA4GH exclude all variant calls within complex, repetitive regions of the genome^25^. In contrast, k-mers inherently capture genetic context, regardless of the structural complexity surrounding them in the genome. Moreover, k-mers are especially robust for evaluating sequences in highly diverged sequences, where mapping based approaches cannot map reads to call variants.

Lastly, Merqury provides an efficient way of determining phase blocks in diploid assemblies. In the past, phase blocks were defined based on heterozygous SNPs, measured by aligning each haplotype to one another^9^ or by mapping to a reference genome^8^. These alignment-based approaches may not consider the full genome when the identity between the two haplotypes is lower than the alignment threshold, or the alignment is confused by genomic repeats. Moreover, a reference genome may not represent the entire haplotype of an assembled individual, thereby omitting haplotype-specific sequences from the analysis. The phase blocks measured by Merqury are generated regardless of the haplotype being assembled, and provide more reliable phasing information for allele-specific studies.

## Methods

### Counting k-mers with Meryl

Meryl is a tool for counting and working with sets of k-mers that was originally developed for use in the Celera Assembler^42^ and has since been adopted and maintained as part of Canu^32^. Here we have improved Meryl’s efficiency and extended it to support a variety of functions useful for k-mer-based assembly validation. A set of k-mers and their associated counts is termed a k-mer database. The count is the number of times a k-mer occurs in some collection of sequences. The k-mer database is stored in sorted order, similar to words in a dictionary. Meryl comprises three modules: one for generating k-mer databases, one for filtering and combining databases, and one for searching databases.

The counting module uses two different algorithms: one for k-mers up to size 16 and the other for k-mers up to size 64. For small k-mers, Meryl directly counts the number of times each k-mer occurs in the input sequences. An array of 4^k^ 16-bit integers is allocated. Each k-mer is converted to an integer index into the array, and the cell for that k-mer is incremented. When any cell exceeds the maximum possible value that can be represented, the width of the array is extended by allocating a supplementary array of 4^k^ bits. For large k-mers, Meryl generates lists of all the k-mers present in the input sequences, sorts each list, then scans each to determine how many times a specific k-mer occurs. Each k-mer is split into a prefix and a suffix. The prefix is used to select a list, and only the suffix is added. A trade-off is made between a small prefix (which would result in a only a few lists, each storing many suffixes) and a large prefix (which would result in many lists). As we do not know how big each list will be, the lists must be able to grow as needed. Each list is therefore an array of memory blocks where each block can store a few thousand k-mers. While counting, the memory usage of the lists is tracked, and if a user-supplied memory limit is reached, the lists are sorted, k-mers are counted, and output written to an intermediate database. After all k-mers are processed, the intermediate, sorted databases are efficiently merged into one.

With one or more databases on disk, Meryl can filter or combine k-mers to create new databases. Each database is stored in 64 independent pieces, and each piece can be processed in parallel. Meryl can filter a database by count (e.g. less-than, greater-than or equal-to some user supplied constant), or by fraction of distinct k-mers in a database (e.g. the most common 5% of the k-mers). It can modify the count of every k-mer in a database by a constant (e.g., add 1, subtract 1, multiply by 2). Meryl can also output the union or intersection of multiple databases, setting the count of a k-mer to the minimum, maximum, sum of all copies of the k-mer or as the count of the first database. It can output the difference of databases (e.g. the k-mer occurs only in the first database) or the symmetric-difference (e.g. the k-mer occurs in exactly one database). Conveniently, any number of these operations can be combined into one command, using a reverse-polish-notation inspired format. The following example invocations are used in Merqury:

1. Write the k-mers that occur in both `db1.meryl` and `db2.meryl` to `db3.meryl`. Set the count of each output k-mer to the sum of the counts in the input k-mers:
`meryl union-sum db1.meryl db2.meryl output db3.meryl`
2. Output k-mers that occur in both `db4.meryl` and `db5.meryl`, additionally requiring the k-mer in `db5.meryl` to be unique. The count of the output k-kmer is set to the count of the k-mer in the first input to the `intersect` operation, namely `db4.meryl`:

~~~
meryl output db6.meryl \
 intersect \
  db4.meryl \
  [equal-to 1 db5.meryl]
~~~
3. For each k-mer in `asm.fasta`, output the (0-based) coordinate of the kmer in the sequence, the forward and reverse k-mer sequences, and the count of the k-mer in

~~~
db7.meryl :
meryl-lookup -dump -sequence asm.fasta -mers db7.meryl
~~~

Meryl includes a C++ API to extend its functionality. For example, random lookups can be added using either the simple existence of a k-mer in a database, or the count associated with a k-mer. On the command line, lookups can return the number of k-mers a sequence shares with a database, a list of each k-mer in a sequence annotated with the count the k-mer has in a database, or a filtered list of input sequences based on the presence or absence of k-mers in the database.

### Evaluating assemblies with Merqury

#### Copy number spectrum (spectra-cn plot)

Given a genome size *G* and tolerable collision rate *p*, an appropriate *k* can be computed as *k* = log_4_ (*G*(1−*p*)/*p*)^43^. However, alterative values of *k* may be desirable for different scenarios, e.g. increasing *k* has the effect of increasing the relative fraction of 1-copy k-mers, which may be useful for genomes with low heterozygosity, but this also increases the fraction of erroneous k-mers. Decreasing *k* reduces the fraction of erroneous k-mers, but increases the fraction of repetitive k-mers. Once an appropriate size if *k* is determined (typically 18∼21), we count the canonical k-mers observed in the assembly and in the accurate, whole-genome read set. A typical k-mer spectrum for a heterozygous diploid genome consists of two primary peaks, one representing k-mers that are 1-copy in the diploid genome (heterozygous, on a single haplotype) and one representing those that are 2-copy in the diploid genome (homozygous, on both haplotypes or two copies on one haplotype) (**Fig. 1a**). The 2-copy k-mers appear with a frequency approximately equal to the average depth of sequencing coverage, where the 1-copy k-mers appear with frequency approximately equal to half the sequencing coverage. If a genome is entirely homozygous, only the 2-copy peak may appear, and if the genome is extremely heterozygous, only the 1-copy peak may appear. With sufficient sequencing coverage (to separate the peaks along the axis), and a proper choice of *k*, both peaks are visible for most genomes. Using the multiplicity of the k-mer counts, and modeling the k-mer survival rate (i.e. how many k-mers are unaffected by sequencing error), it is possible to predict the size and repeat content of a genome from the k-mer spectrum alone^35^.

The spectra-cn plot was introduced by Mapleson *et al*.^17^, which colors k-mers of the read set by their copy numbers in the assembly. In addition to the original stacked version of the spectra-cn plot (**Fig. 1b**), we provide additional options to plot the unstacked copy number spectrum (**Fig. 1c**). We have found this style more useful for visually detecting abnormal k-mer copy numbers and their distribution in an assembly.

#### Assembly spectrum (spectra-asm plot)

Similar to the spectra-cn analysis, we can color each k-mer in the read set by the assembly in which it is found. This becomes useful when two haploid assemblies are evaluated. This way, we can detect k-mers that are present only in one assembly, k-mers shared in both assemblies, and k-mers not present in the assembly and only found in the read set (**Fig. 2**).

#### K-mer completeness

We define a “reliable k-mer” as a k-mer that is truly in the genome and unlikely to be caused by sequencing error. With exact k-mer counts, it is easy to filter out low-copy k-mers that are likely to represent sequencing errors. We use the same strategy as Koren *et al*.^10^ to find the cutoff. In brief, we take the histogram of the k-mer counts and set the multiplicity (number of times we see the k-mer in the read set) as *x* and counts (number of k-mers with *x* multiplicity) as *y*. When differentiating the histogram, we compute the slopes and the first k-mer multiplicity with a positive slope defines the reliable k-mer threshold. Examples of these cutoffs are shown as dashed lines in **Fig. 3c-d**. The k-mer completeness is calculated as the fraction of reliable k-mers in the read set that are also found in the assembly. For repetitive genomes, erroneous read set k-mers can sometimes appear above this threshold due to recurring errors in high-copy repeat families, but this is a rare.

#### Consensus quality (QV) estimation

We can also use k-mers to estimate the frequency of consensus errors in the assembly. We use a binomial model of k-mer survival, and assume all k-mers in the assembly should be found at least once in the read set. Here, we use the containment score from Mash Screen^38^ to estimate consensus accuracy. In brief, we estimate the probability *P* that a base in the assembly is correct as:

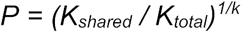

Where the *K*_*total*_ is the total number of k-mers found in an assembly and *K*_*shared*_ are the number of shared k-mers between the assembly and the read set. If the read set is assumed to completely cover the genome, any k-mer found only in the assembly (*K*_*asm*_ *= K*_*total*_ *-K*_*shared*_) likely reflects a base error in the assembly consensus. Hence, the error rate *E* can be defined as:

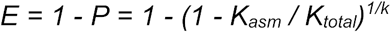

Using this formula, the widely used Phred^44^ quality score (often denoted as QV) can be computed by treating the *E* as base error probability:

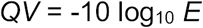

Note that this QV estimate depends on the coverage and quality of the read set. The assembly QV will be underestimated if the read set does not contain all true k-mers of the genome due to low or heavily biased sequencing coverage. Conversely, the QV may be overestimated if the read set contains a high proportion of false k-mers (e.g. due to the combination of extreme coverage and sequencing error). Lastly, the k-mer survival model assumes both k-mers and errors are independent, but k-mers overlap one another and errors tend to cluster together. As a result, choice of *k* also affects the QV estimates, with *k* ≈ 21 recommended by Mash based on empirical testing.

#### Positions of k-mers for mis-assembly detection

Merqury reports the positions of the k-mers found only in an assembly for further investigation in .bed and .tdf formats that can be loaded into most genome browsers. In addition, the k-mers found in unexpected copy numbers (i.e. false duplications) are also provided as .bed and .tdf files. The format details are described at:

https://genome.ucsc.edu/FAQ/FAQformat.html#format1

https://github.com/igvteam/igv/wiki/TDF-Format

### Evaluating phasing completeness with parental genomes

#### Haplotype-specific k-mers (hap-mers)

Parental haplotype markers can be obtained directly from the parental or ancestral genomes^10^. In brief, distinct k-mers found in only one parent are collected, and the erroneous low-frequency k-mers are filtered out. This filtering strategy relies on the k-mer count histogram, where the cutoff for identifying reliable k-mers is computed as described above. When the child’s short-read data is also available, the inherited haplotype-specific markers can be obtained by intersecting the child’s k-mers with the parental marker sets. This time, we keep the k-mer counts from the child’s reads for further ploidy estimation (**Fig 3**).

#### Hap-mer blob plot

To get a global visual representation of the phasing completeness on assembled sequences, we can count the total number of hap-mers found on each contig or scaffold (**Fig 4c-d**). Here, each axis becomes the number of hap-mers found in a sequence entry (contig or scaffold). Each circle (blob) represents a sequence, the size being relative to the length of the sequence. Sequence bubbles near the diagonal represent mixtures of markers from both haplotypes, while bubbles closer to a haplotype axis are predominately comprised of markers from a single haplotype.

#### Phase block statistics and switch error

Whenever a hap-mer switch occurs, Merqury can flag that position in the assembly and output a haplotype block report. This feature is useful for identifying phase blocks from a partially or completely phased assembly such as FALCON-Unzip^9^, FALCON-Phase^12^, or Supernova2^4^. Merqury defines a phase block as a continuous sequence with at least two hap-mers originating from the same haplotype (**Fig. 4a**). Usually, short-range switches are caused by consensus, rather than phasing, errors. By default, Merqury allows at most 100 hap-mer switches per 20 kbp as short range switches within a phase block. Each unexpected hap-mer found will be counted as a switch error, but will not necessarily terminate the phase block. Ideally, when no switches are found, the phase block N50 will be the same as the scaffold N50 and the sum of the phase blocks will be identical to the assembled sequences. In reality, a scaffold often does not end with a haplotype specific sequence, so the total phase block length is shorter. Trio-binned or haplotype-resolved assemblies are a special case, where the entire haplotype assembly is essentially a single block. Still, in this case, hap-mers from the other haplotype can be counted as switch errors. Merqury also provides an option to restrict phase blocks to contigs and break the blocks at any gap. At the end, Merqury reports the number of switches and total hap-mers on each block along with the switch error rate in order to identify blocks with more frequent switch errors.

#### Hap-mer copy number spectrum

The total k-mer spectrum of the assembled individual is also useful for tracking the fates of haplotype-specific k-mers in a diploid assembly. Similar to the overall copy number analysis performed with spectra-cn plots, we can count k-mers in each haplotype assembly and estimate the completeness of haplotype-specific assembled bases compared to the hap-mer sets. For example, by plotting separate histograms for hap-mers of different copy numbers in the assembly (**Fig 6**), we can identify whether the assembly is artificially collapsing or duplicating sequence in each haplotype. If hap-mers appear over (or under) represented by the assembly relative to the read set, it is an indicator of artifactual duplication (or absence) of haplotype-specific sequence. When evaluating a pseudo-haplotype assembly, which is designed to collapse or pick one haplotype as much as possible, we can count the number of hap-mers present in the child’s read set but not present in the pseudo-haplotype assembly and use this to quantify the amount of under-represented haplotype sequence. These missing hap-mers could then be used to identify a set of alternative haplotype reads that were incorrectly excluded from the assembly.

### Assemblies

All TrioCanu assemblies were downloaded from Koren *et al*.^10^, available at https://obj.umiacs.umd.edu/marbl_publications/triobinning/index.html. The *A. thaliana* F1 FALCON-Unzip assembly was obtained from Chin *et al*^9^. We generated a Canu assembly to show a typical example of an interim mixed-haplotype assembly that has not been polished or purged of haplotypic duplications. The same Pacific Bioscience reads was used for all three assemblies, obtained from Chin *et al*^9^.

The Canu assembly was generated with Canu 1.9 release version using the following command:

~~~
canu -p canu -d athalF1_notrio genomeSize=130m ‘corOutCoverage=100’ ‘batOptions=-dg 6 -db 6 -dr 1 -ca 500 -cp 50’ ‘batMemory=200’ -pacbio-raw *.fastq.gz
~~~

### Haplotype specific k-mers (hap-mers) for *A. thaliana F1* and NA12878

Appropriate size of k was obtained as k=18 for the *A. thaliana* F1 with 130∼260 Mbp genome size and k=21 for NA12878 with 3.2∼6.4 Gbp genome size using `$MERQURY/best_k.sh`.

As parental Illumina sequencing was not available for this F1, the parental genome assemblies from Chin *et al*^9^. were used to obtain parental specific k-mers. Each assembly from the inbred Col-0 and Cvi-0 lines were downloaded from:

https://downloads.pacbcloud.com/public/dataset/PhasedDiploidAsmPaperData/FUNZIP-PhasedDiploidAssemblies.tgz.

Meryl databases for the parental strains were built directly with `meryl count k=18 output $hap.meryl $hap.fasta` for each haplotype assembly.

The parental Illumina whole-genome sequencing sets for NA12878 were downloaded from the high coverage dataset of the 1000 Genomes Project (NA12891 and NA12892) and combined with Illumina Platinum Genomes Project data from PRJEB3381. Illumina whole-genome sequencing of NA12878 was downloaded from PRJEB3381.

All Meryl databases from sequencing read sets were built with `$MERQURY/_submit_build.sh`. Once the k-mer databases were built, inherited hap-mers were obtained with `$MERQURY/trio/hapmers.sh.`

### Merqury on all assemblies

Merqury was run for the *A. thaliana* F1 and NA12878 with the following command line for diploid assemblies, where `$hap1` and `$hap2` are maternal (mat) and paternal (pat) for the TrioCanu assemblies, and primary (pri) and alternate (alt) for FALCON-Unzip assemblies.

~~~
$MERQURY/_submit_merqury.sh $sample.k21.meryl mat.inherited.meryl pat.inherited.meryl $hap1.fasta $hap2.fasta $asm_name
~~~

The *A. thaliana* F1 Canu assembly was run with:

~~~
$MERQURY/_submit_merqury.sh $sample.k21.meryl mat.inherited.meryl pat.inherited.meryl canu.contigs.fasta Canu
~~~

### BUSCO

BUSCO v3 was run on using `embryophyta_odb9` for the *A. thaliana* Canu assembly and the combined Col and Cvi TrioCanu assembly with the following commands:

~~~
python run_BUSCO.py -i asm.fasta -o SAMPLE -l embryophyta_odb9 -m genome -c 16 -sp arabidopsis
~~~

For NA12878, BUSCO was run in the same way, using `mammalia_odb9` for the combined TrioCanu maternal and paternal assembly. BUSCO scores for each haplotype of *A. thaliana F1* and NA12878 were obtained from Koren et al.^10^ Supplementary Table 2.

### QV estimates

CPU Time, memory consumption, and disk usage was measured for generating QVs on each haploid assemblies and the combined diploid assembly. A Intel(R) Xeon(R) Gold 6140 CPU @ 2.30GHz node was used allowing up to 24 CPUs. Detailed node information is available at https://hpc.nih.gov.

#### Meryl-based QV

Meryl-based QV estimation to benchmark computing resources was evaluated for the counting (count), merging (union-sum), and QV steps with the following command:

~~~
$MERQURY/eval/qv.sh NA12878.k21.meryl mat.fasta pat.fasta meryl_qv
~~~

This generates Meryl databases for mat.fasta and pat.fasta, does a union for the databases, and generates QV scores for all three combinations (maternal, paternal, and both).

#### Mash-based QV

Mash-based QV estimation was performed using sketch size of `1000000` with the same k-mer size of 21:

~~~
mash sketch -s 1000000 -k $k $asm
mash screen -p $cpus $asm.msh ‘cat $input_fofn | tr ‘\n’ ‘‘’ >
$name.msh.idy
cat $name.msh.idy | awk -v name=$name ‘{print name”\t”$2”\t”-
10*log(1-$1)/log(10)”\t”(1-$1)}’ | tr ‘/’ ‘\t’ > $name.msh.qv
~~~

#### Mapping-based QV estimates

The Illumina WGS reads used to build the Meryl database were aligned to both mat.fasta and pat.fasta using BWA^45^. Base pair errors were called using FreeBayes^46^ v1.3.1 `--skip-coverage 600`, skipping variant calling in regions with >600x read depth to help prevent unnecessary computing on high coverage regions that violate the diploid assumption. The exact commands used were:

~~~
# Add mat and pat at sequence names to prevent naming collisions in contigs
sed ‘s/>/>mat_h_/g’ mom.fasta > mat.fasta
sed ‘s/>/>pat_h_/g’ dad.fasta > pat.fasta
# bwa indexing
bwa index both.fasta
# bwa alignment
bwa mem -t 24 both.fasta F_1.fastq.gz F_2.fastq.gz > na12878.sam
# Sorting and converting to bam
samtools sort -@24 -O bam -o na12878.bam -T na12878.tmp na12878.sam
# Freebayes variant calling
freebayes --bam na12878.bam --skip-coverage 600 -f both.fasta |
bcftools view --no-version -Ou > na12878.tmp.bcf
bcftools index na12878.tmp.bcf
# Normalize indels
bcftools view -Ou -e’type=“ref”’ na12878.tmp.bcf | bcftools norm -
Ob -f both.fasta -o na12878.bcf --threads 24 bcftools index na12878.bcf
# Filter out low quality variant calls
bcftools view -i ‘QUAL>1 && (GT=“AA” || GT=“Aa”)’ -Oz --threads=24 na12878.bcf > na12878.changes.vcf.gz
bcftools index na12878.changes.vcf.gz
# Get number of bases called as errors
bcftools view -H -i ‘QUAL>1 && (GT=“AA” || GT=“Aa”)’ -Ov na12878.changes.vcf.gz | awk -F “\t” ‘{print $4”\t”$5}’ | awk ‘{lenA=length($1); lenB=length($2); if (lenA < lenB) {sum+=lenB-lenA} else if (lenA > lenB) {sum+=lenA-lenB} else {sum+=lenA}}
END {print sum}’ > na12878.numvar
# Get number of bases with alignment
samtools view -F 0×100 -u na12878.bam | bedtools genomecov -ibam -
-split > aligned.genomecov
awk -v l=3 -v h=600 ‘{if ($1==“genome” && $2>l && $2<h) {numbp +=
$3}} END {print numbp}’ aligned.genomecov > na12878.numbp
# QV calculation
NUM_BP=‘cat na12878.numbp‘
NUM_VAR=‘cat na12878.numvar‘
QV=‘echo “$
NUM_VAR $NUM_BP” | awk ‘{print (−
10*log($1/$2)/log(10))}’‘
echo $QV
~~~

This pipeline is used in the Vertebrate Genomes Project, and the code used is available from https://github.com/VGP/vgp-assembly/tree/master/pipeline/ under the “bwa”, “freebayes-polish”, and “qv” directories.

## Declarations

### Ethics approval and consent to participate

Not applicable.

### Consent for publication

Not applicable.

### Availability of data and materials

Merqury is openly available from https://github.com/marbl/merqury. Meryl is available from https://github.com/marbl/meryl. Meryl v1.0 release was used in this manuscript. Downloadable links to the Meryl k-mer databases of the assembled individuals and hap-mers are available for *A. thaliana* and NA12878 on the Merqury GitHub page.

## Authors’ contributions

A.R. conceived the project and implemented Merqury. B.P.W. designed and implemented Meryl. A.R., B.P.W., S.K., and A.M.P. designed the methods. A.R. and S.K. performed analyses. A.R. and A.M.P. coordinated the project. All of the authors wrote the manuscript and approved the final manuscript.

## Acknowledgements

The authors would like to thank the members of the Vertebrate Genomes Project assembly working group for helpful discussions, including Marcela Uliano da Silva and Giulio Formenti. We also would like to thank Bernardo Clavijo of the Earlham Institute for his very insightful comments on a preprint of this manuscript. This research utilized the computational resources of the NIH HPC Biowulf cluster (https://hpc.nih.gov).

